# Comparing neurocognitive mechanisms of mathematical ability and fluency in children: insights from an fNIRS study

**DOI:** 10.1101/2024.11.15.623735

**Authors:** Alethea Yen Ning Yeo, Fengjuan Wang, Azilawati Jamaludin

## Abstract

**Background:** Early proficiency in mathematics is a strong predictor of later academic success and life achievement, considering the practical skills that mastering the subject enables students to equip. Yet, there exists a paucity of research into the neural mechanisms supporting mathematical abilities in young children. Recent research utilises resting state functional connectivity (RSFC), a measure of the coherence of brain activity among brain regions in the absence of tasks, to understand the functional roles of these regions.

**Methods:** We analysed the RSFC of 45 children to investigate the intrinsic cognitive processes underpinning arithmetic processing in three regions of interest (ROIs): middle frontal gyrus, inferior parietal lobule, and precuneus. Correlations between RSFC among these regions and mathematics or math fluency scores, derived from the Wechsler Individual Achievement Test (WIAT-III), were examined.

**Results:** RSFC between the right precuneus and both the ipsilateral middle frontal gyrus and inferior parietal region may be associated with arithmetic processing speed and accuracy, while cross-hemispheric RSFC between the right precuneus and the left inferior parietal lobule appears to be associated with problem-solving and numeracy skills. RSFC between the right precuneus and left inferior parietal lobule differed in children performing below the 10^th^ percentile in mathematics (out of 45 participants).

**Conclusions:** The results suggest that children of the same age may follow different neural development trajectories. More targeted and differentiated interventions are essential to offer additional and early support for students struggling with mathematics.

## 1. Introduction

Early numerical competency is an important predictor of academic success and life achievement, particularly in maintaining health, employment, and even avoiding incarceration [[1],[2]]. Longitudinal studies show that school-entry mathematical skills have the greatest predictive power for later academic performance, over other cognitive skills such as literacy [2]. Morgan, Farkas and Wu [3] showed that 70 percent of children who had ended kindergarten below the 10^th^ percentile in mathematics remained below the 10^th^ percentile in fifth grade. Consequently, recent research has focused on exploring the factors that influence the development of this skill from an early age. Naturally, this leads to the discussion on the interplay between innate abilities and environmental influences. While it is essential to recognise the impact of the environment on a child’s early numeracy skills such as parental engagement in mathematics learning, we posit that the specific neurocognitive mechanisms supporting early mathematical skills also play a critical role.

The human brain is a complex organ, characterised as a structurally or functionally interconnected network which coordinates and processes a continuous flow of information [[4],[5]] to support brain-behaviour mechanisms. Structurally, the brain is organised into specialised regions which work together to process neural signals and facilitate perception and cognition [6]. Where brain regions are more activated than others, a higher hemodynamic response would be detected through imaging techniques like fNIRS [7]. Functionally, beyond quantifying neural activity during tasks, research has employed resting state functional connectivity (RSFC), a measure of the temporal synchronisation of brain activity among anatomically separated brain regions, in the absence of tasks [8]. Such activity is known as spontaneous brain activity which reflects the fluctuations in brain activity during resting state. At this natural state, there is neither overt perceptual input nor behavioural output. As this state may be achieved without active participation from subjects, it serves as a crucial experimental paradigm for studying brain function across diverse participants, from young children to clinical patients [[9],[10]]. During specific functional tasks, brain regions that co-activate remain functionally correlated during resting states, preserving task-related specificity. There is a strong relationship between RSFC and task-evoked functional connectivity, whereby findings from RSFC largely overlap findings that would have been observed with a specific task paradigm [[8],[11],[12]]. While some studies show differences in functional connectivity in task-based versus resting state conditions [[13],[14]], others have shown that the brain’s functional network organisation during task is shaped primarily by the intrinsic network organisation during rest [11]. Importantly, individual differences in functional connectivity are preserved during both task and resting conditions [14]. RSFC has also been used to demonstrate group-specific features [15], such as differences in connectivity between mathematicians and non-mathematicians [16]. In addition, RSFC is useful in elucidating potential biomarkers associated with atypical learning development [17]. While these findings were garnered from fMRI studies, the insights gleaned regarding resting state and RSFC are nonetheless relevant.

The development of mathematical proficiency relies on distinct yet complementary neurocognitive processing systems [18]. These include systems promoting symbolic numbers perception (e.g., Arabic numerals), memory, and cognitive control processes that manipulate representations of quantity [19]. Core systems of numerical cognition involve the quantity representation system in the intraparietal sulcus [[20],[21]] and the visual number form processing system in the ventral temporal-occipital cortex [[22],[23]]. These regions cooperate to produce semantic representations of quantity and enable efficient manipulation of numbers for problem-solving [24]. For higher-order mathematical learning, these regions work with memory systems that maintain and manipulate quantities. Working memory is regulated by the visuospatial attention system and cognitive control system, which is also an integration hub of various functional circuits engaged in problem-solving [[25],[26],[27],[28]]. Developmental studies show from childhood to adulthood, activation in the frontal regions decreases, while activation in the parietal regions increases [[29],[30],[31],[32]]. This frontal to parietal shift reflects a shift from reliance on memory and cognitive control systems to specialised functional networks. This is consistent with the interactive specialisation framework, suggesting cognitive development depends on selective strengthening and weakening of neuronal networks to produce specialised and interconnected functional modules with time [33]. Freeing the frontal cortex from computational load makes available valuable processing resources for complex problem-solving, which is important to improve mathematical competency [34]. Accordingly, early mathematical problem-solving skills progress from laboured strategies such as counting to more efficient direct retrieval of mathematical facts [35].

Mathematical proficiency is further described as a concordance of various brain regions and networks that vary along a functional continuum, with domain-general to domain-specific functions [36]. Domain-general functions, such as working memory and visuospatial reasoning, support general learning and information processing [37]. Domain-specific functions involve quantitative processing which is less relevant in other learning domains like reading. This includes numerical representation and symbolic knowledge about number orders [38]. Importantly, brain imaging studies reveal cortical regions are not devoted to a specific task, but rather contribute to a variety of domain-specific and domain-general functions [39]. This study focuses on numeracy and math fluency—two components of mathematical proficiency. Arguably, numeracy involves domain-specific brain functions for understanding and applying mathematical concepts, while math fluency recruits domain-general functions for arithmetic fact retrieval and quick manipulation of numbers mentally. There is consensus that mathematical learning can be represented as a pyramid, in order of fluency at the base, mathematical reasoning, and problem solving at the apex. As fluency allows one to quickly recall mathematical facts and concepts, spot patterns and make generalisations, more complex schemas of reasoning can be built upon [40]. However, some posit that mathematical reasoning can occur without arithmetic fluency. This implies the ability to reason relatively without quantitative evidence [41]. With these conflicting theories, studies into the neural distinction between these aspects of mathematical ability would be critical as neural indicators such as RSFC are more objective and may provide insight on mathematical learning.

Applying a seed-based approach, studies have investigated the neural differences supporting mathematical skills in the regions: middle frontal gyrus, inferior parietal lobule, and precuneus. The middle frontal gyrus, functionally identified as the dorsolateral prefrontal cortex (dlPFC; Brodmann’s area 46), plays a critical role in supporting working memory through the processing of visuospatial information and the maintenance of mental representations [[42],[43]]. In a task-based study by Rivera et al. [31], participants aged eight to 19 evaluated the accuracy of arithmetic equations. Younger participants had a greater activation in the dlPFC, indicating heightened demands on working memory and attention. Notably, with adults, mathematicians yielded a focal dlPFC activation, unlike in non-mathematicians where a low level of expertise was reflected by diffuse activity across broader brain regions (e.g., inferior frontal gyrus and right inferior parietal lobule) [44]. The dlPFC also supports proactive control, which tends to increase with age [45] and is associated with increased working memory and mathematical proficiency in children [[46],[47]]. A task-based fNIRS study found that greater proactive engagement in third-graders was related to increased left lateral prefrontal cortex activation during reactive beneficiary situations and improved mathematical performance [48]. Further, as elementary mathematics training coincides with prefrontal cortex development [49], understanding the activity within this region and its interactions with parietal regions will be crucial for elucidating variations in mathematical proficiency among school-age children.

Next, the inferior parietal lobule comprises the intraparietal sulcus, angular gyrus, and the supramarginal gyrus. The intraparietal sulcus supports numeracy in typically developing individuals [50], and is structurally and functionally different in individuals with learning disabilities, such as developmental dyscalculia, a neural deficit in number processing and arithmetic [[51],[52]]. Evans et al. [53] showed RSFC between the intraparietal sulcus and prefrontal cortex predicted gains in mathematical proficiency over six years. Similarly, Jolles et al. [54] reported that increased intraparietal sulcus RSFC with other regions such as the lateral prefrontal cortex, correlated with individual performance gains in arithmetic tutoring for third-graders. Moreover, hyperconnectivity between the intraparietal sulcus and bilateral frontoparietal regions was observed in children with mathematical learning difficulties [55]. These results highlight the behavioural significance of plasticity in intraparietal sulcus circuits and suggest the potential of using RSFC among these regions as biomarkers for arithmetic proficiency. However, while understanding which regions predict gains in arithmetic ability is useful to identify students likely to perform poorly in mathematics in future based on current neuroimaging data, gains in arithmetic ability will often be inherently confounded by other environmental factors, such as changing mathematics curriculums. Therefore, while these neural mechanisms predict learning gains, they may not accurately reflect underlying neural mechanisms of specific mathematical processes. Further research could explore RSFC’s association with specific mathematical domains, and whether the same regions are involved across domains. Arithmetic training shifts activation from the intraparietal sulcus areas involved in magnitude processing, to regions that support arithmetic fact retrieval, such as the angular gyrus (BA 39) [[56],[57],[58]]. Crucially, a study by Wu and colleagues [59] showed that the angular, but not supramarginal gyrus or intraparietal sulcus, responded differently to task difficulty in mental arithmetic. This demonstrates the unique contribution of the angular gyrus in handling increased cognitive demands in mathematical tasks. This may be explained by the symbol-referent mapping hypothesis which suggests that the left angular gyrus mediates the mapping of arithmetic problems onto solutions stored in memory [60]. Therefore, activation of this region, which increases with arithmetic training [61], increases when solving simpler problems and during problem-solving through fact retrieval rather than calculation [[62],[63]]. Finally, the supramarginal gyrus (BA 40) is strongly engaged during numerical problem-solving tasks that require active storage and manipulation of items in the working memory [25].

The precuneus (BA 7) is a highly interconnected associative hub within the posterior parietal lobule, facilitating tasks such as episodic memory retrieval, visuospatial imagery and self-referential processing [[64],[65],[66]]. Given its role in integrating information from frontal and parietal regions, the precuneus is thought to subserve higher-order mathematical reasoning and problem-solving. During calculation, children engage regions including the bilateral precuneus [19], which predicts children’s composite mathematical skills during symbolic number comparison [67]. Despite the importance of the precuneus as a mediator among frontoparietal regions important for mathematical processing, few studies have explored the relationship between the associations of the precuneus and other regions, with mathematical domains, such as problem-solving, in children.

The angular gyrus, supramarginal gyrus and intraparietal sulcus are distinct regions and ideally, should not be conflated into one region. However, while fNIRS provides better spatial resolution [68] than other modalities such as EEG, the precise locations of small neighbouring regions such as the intraparietal sulcus and angular gyrus cannot be as accurately demarcated as modalities like fMRI can, which most existing studies have been based on. This is especially so in children with smaller brains and narrower distinctions between regions. Hence, we investigated patterns in cross-regional functional connectivity with less specific but more anatomically distinct regions.

### 1.1 Current study

In the current study, we aim to clarify how neural mechanisms in frontoparietal regions differ in mathematical problem-solving and math fluency, among school-entry children. While these regions may share overlapping functions, such as supporting arithmetic fact retrieval and logical reasoning, the distinct domains of mathematics and math fluency are often studied separately. Thus, distilling the similarities and differences in neural mechanisms supporting each domain can provide a nuanced understanding of the neurobiological basis of mathematical proficiency. Secondly, we ask if the RSFC among regions associated with mathematics or math fluency would be significantly different in children below the 10^th^ percentile. By investigating this question, we probe into the neural underpinnings of numerical competency, which may reveal common weaknesses in the low performing children in each domain. This would be crucial for developing neurobiologically informed education strategies.

We predicted that the neural mechanisms underlying mathematical problem-solving and math fluency will differ. Problem-solving may require a stronger connectivity between the precuneus and both the middle frontal gyrus and the inferior parietal lobule. This is due to the cognitive demands of logical reasoning, visuospatial processing, attention, and working memory manipulation. Conversely, math fluency may rely on stronger connectivity within the posterior parietal regions (precuneus and inferior parietal lobules) to facilitate arithmetic fact retrieval. Additionally, we hypothesised that RSFC among the regions associated with mathematics or math fluency will be different in children below the 10^th^ percentile. Specifically, we predicted that the frontoparietal RSFCs will be significantly different in these children, for both mathematical domains. This is due to the well-established shift from frontal to parietal regions with increasing mathematical competency and development [69].

## 2. Materials and methods

### 2.1 Participants

This study comprised a random sample of 45 children (age range: 6.17 to 7.92 years old; M_age_ = 6.96 years) from five primary schools across Singapore. The sample comprised 46.7% females and 53.3% males. Informed written consent was obtained from parents/guardians on behalf of the children, prior to participation in accordance with the Institutional Review Board of the Research Integrity and Ethics Office of Nanyang Technological University (IRB-2017-10-030 and IRB-2019-07-042). None had any known neurological or learning disorders.

### 2.2 Data collection

#### 2.2.1 Wechsler Individual Achievement Test—Third Edition (WIAT-III)

WIAT-III^72^ is a standardised academic achievement test that assesses a child’s educational progress in areas such as Mathematics. This study utilised results from the Mathematics and Math Fluency domains. The Mathematics domain included Problem-solving and Numerical Operations subtests. In the latter, children solved orally-presented word problems. The questions may require several steps and may be related to money, measurement, geometry, time, or interpreting graphs. To ensure the task was a measure of mathematical knowledge instead of working memory, the children were given paper and pencil. Numerical Operations evaluated calculation skills with basic operations, such as “+” and “-“. Math Fluency, assessed through timed Addition and Subtraction tasks, evaluated calculation speed and accuracy. Standardised scores by age for each subtest were used to derive composite scores for Mathematics and Math Fluency achievements.

#### 2.2.2 NIRS Acquisition

The non-invasive optical imaging tool, functional near-infrared spectroscopy (fNIRS) was used in this study. By projecting near-infrared light (650-950nm [71]) through the skull, fNIRS can quantify the light intensity refracted off the cortical surface into nearby detectors [72]. This technique is used to quantify relative changes in oxyhemoglobin (HbO_2_) and deoxyhemoglobin (HbR) concentrations, as changes in oxygenation of a local cortical area provide indications of neuronal activation [7]. Importantly, fNIRS offers numerous advantages over other imaging modalities. These include portability and affordability, better temporal resolution [[73],[74]], and enhanced tolerance to movements during measurements [75], making fNIRS the most suitable technique for assessing neural activity of children. Further, as fNIRS is portable and compact, it can be used in natural settings such as schools [76], providing stronger ecological validity [77]. However, fNIRS has limited sensitivity to hemodynamic changes in subcortical regions (limited penetration depth of 1.5-2 cm [7]).

Data was recorded at 8.14 Hz with a continuous wave functional NIRS system (NIRSport2, NIRx Medical Technologies LLC, Glen Head, NY, United States). The fNIRS cap contained 15 LED sources and 14 detectors (39 source-detector pairs; inter-probe distance of 3 cm), with probe placement following the international 10-20 standard system (frontal and parietal regions; Fig. 2). During the resting state data collection, participants were given toys for eight minutes [[78],[79]] of free play. Doing so has shown to have good construct validity and test-retest reliability [80]. To ensure there were no carry-over effects from the WIAT-III assessments to the resting state data collection which may have confounded the RSFC data, fNIRS data was collected prior to any WIAT-III assessments.

### 2.3 Analysis methods

#### 2.3.1 fNIRS Data Preprocessing

The resting state fNIRS data was first visually inspected, then preprocessed in NIRS-KIT v3.0 [81] using MATLAB R2023b (MathWorks Inc., Natick, USA) (Fig. 1). Data preprocessing included time point trimming, detrending, motion correction and bandpass filtering to remove noise from data acquisition and physiological sources like respiration. Channels with a signal-to-noise (SNR) ratio less than 2 were excluded. The changes in absorbed NIR-light were subsequently transformed into relative HbO_2_ and HbR levels using the modified Beer-Lambert Law [82], although only the HbO_2_ data was used in the current analyses.

**Fig. 1.**
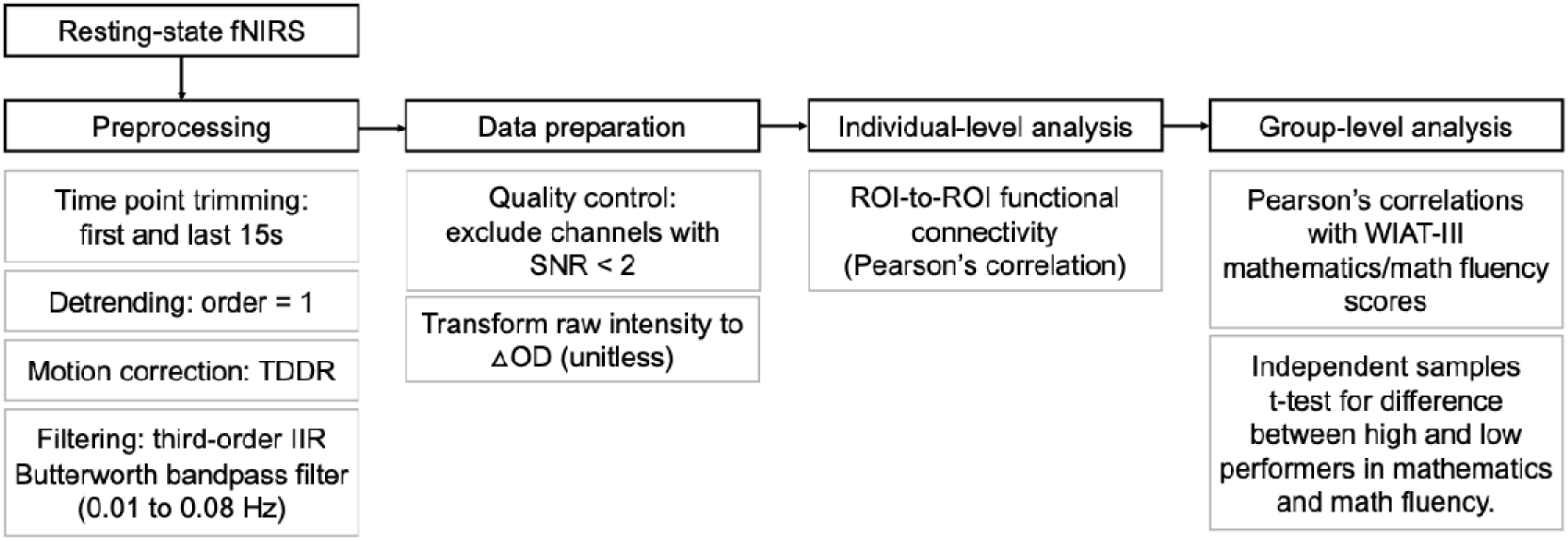
fNIRS data processing pipeline. TDDR, temporal derivative distribution repair; IIR, infinite impulse response filter; SNR, signal-to-noise ratio; OD, optical density.

#### 2.3.2 ROI Selection

Referencing Fig. 2, channels measuring brain activity in the three regions were selected; these included channels covering the dlPFC at the middle frontal gyrus, precuneus and inferior parietal lobule (Appendix A). Selected channels were cross-checked using the MATLAB toolbox, fNIRS optode location decider (fOLD), which more precisely guides the selection of channel positions for fNIRS experiments [83]. This is a traditional and widely accepted approach that defines fiducial points on the scalp according to anatomical landmarks, based on the international 10-20 standard system [84]. Furthermore, multiple studies have shown that there is a reasonable positional correlation between the fiducial points and the anatomical structure of the cerebral cortex [[85],[86]].

**Fig. 2.**
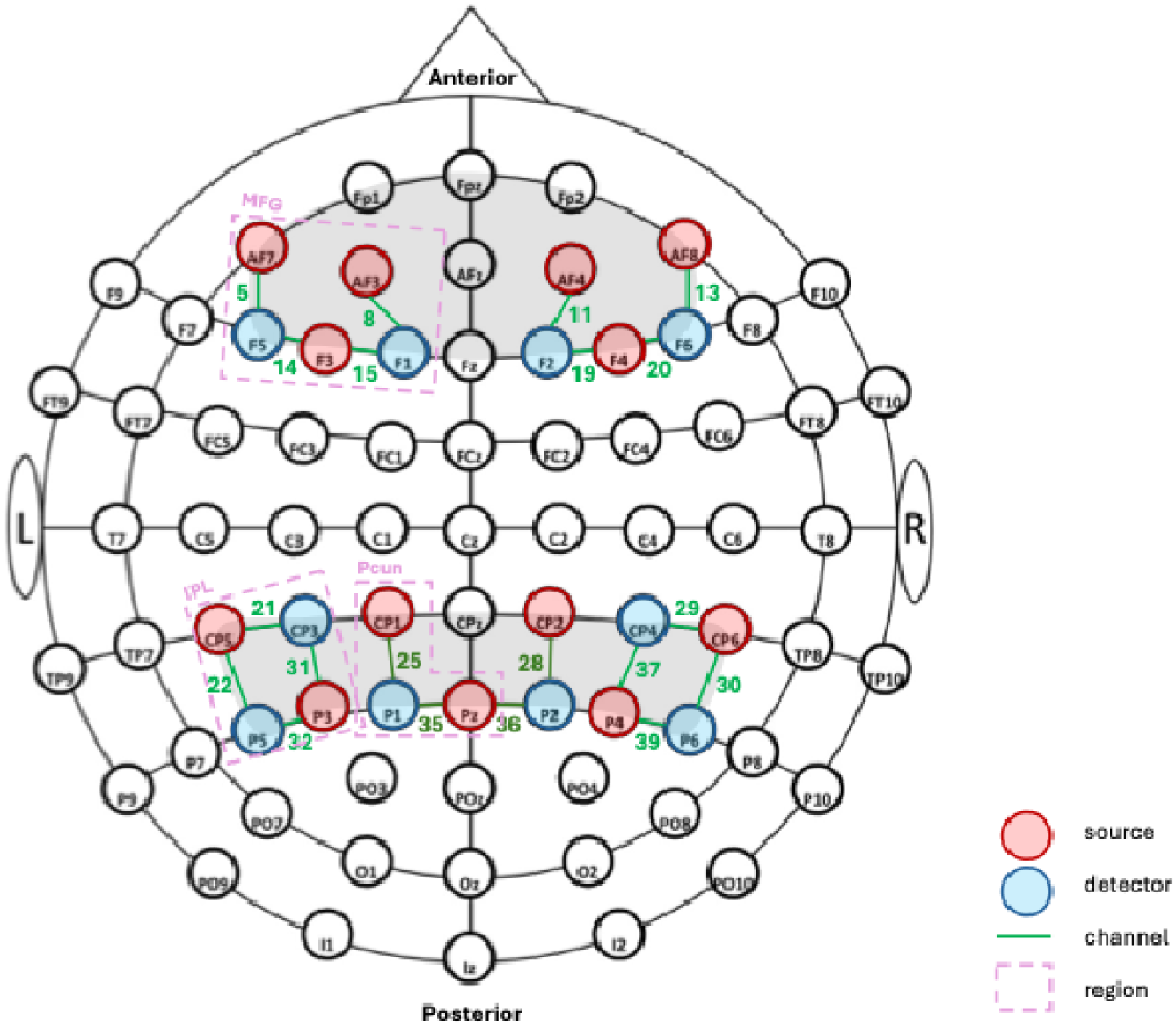
Sources, detectors and channels on the fNIRS cap (shaded grey), in the international 10-20 head space. MFG, middle frontal gyrus; IPL, inferior parietal lobule; Pcun, precuneus.

#### 2.3.3 Statistical analyses

ROI-to-ROI RSFC between two of the three selected regions were correlated with mathematics or math fluency scores. Functional connectivity, assessed from the inter-regional correlations of brain activity, has often been correlated with behavioural measures such as mathematics proficiency, to provide important insights into the functional integration of the brain in relation to mathematical performance [87]. By separately correlating RSFC with mathematics versus math fluency, any differences in the functional integration of the regions in each domain may be nuanced.

Independent samples t-test was used to investigate if ROI-to-ROI RSFC of high performing (HP) children differed from that of lower performing (LP) children. LP children performed below the 10^th^ percentile of the study’s sample. As it was later found that the HP in mathematics were not necessarily HP in math fluency too, this categorisation was done with the mathematics and math fluency scores separately.

## 3. Results

### 3.1 Mathematics and math fluency scores

The means, standard deviations and Pearson’s correlations of mathematics and math fluency scores are shown in Table 1. There was a strong correlation between mathematics and math fluency scores which was expected due to their inherent relatedness in skill. A further paired t-test (Table 2) conducted assured a significant difference between mathematics and math fluency scores. This acts as a control for between-subject differences, as subjects may perform generally better on any math-related task due to their overall arithmetic aptitude.

**Table 1.** Means, Standard Deviations and Pearson’s Correlations between Mathematics and Math Fluency.

| Variables | *N* | *M* | *SD* | Pearson's Correlations |  |
| --- | --- | --- | --- | --- | --- |
|  |  |  |  | 1 | 2 |
| 1. Mathematics | 45 | 105.57 | 15.25 | - | - |
| 2. Math Fluency | 45 | 110.22 | 15.02 | .78** | - |

Note. \*\* $p < .01$

**Table 2.** Paired samples t-test of differences between Mathematics scores and Math Fluency scores for the 45 participants.

| Pair | Paired samples *t*-test |  |  |  |
| --- | --- | --- | --- | --- |
|  | *t* | Sig. (2-tailed) | Cohen's *d* | 95% CI |
| Mathematics –<br>Math Fluency | -3.10 | .003** | -0.46 (modest) | [ -7.66, -1.62 ] |

Note. \*\* $p < .01$

### 3.2 Relating RSFC and mathematics ability

The RSFC of each pair of ROIs were correlated against mathematics or math fluency scores (Appendix B), with the four ROI pairs showing a significant correlation with mathematics and/or math fluency scores illustrated in Fig. 3.

**Fig. 3.**
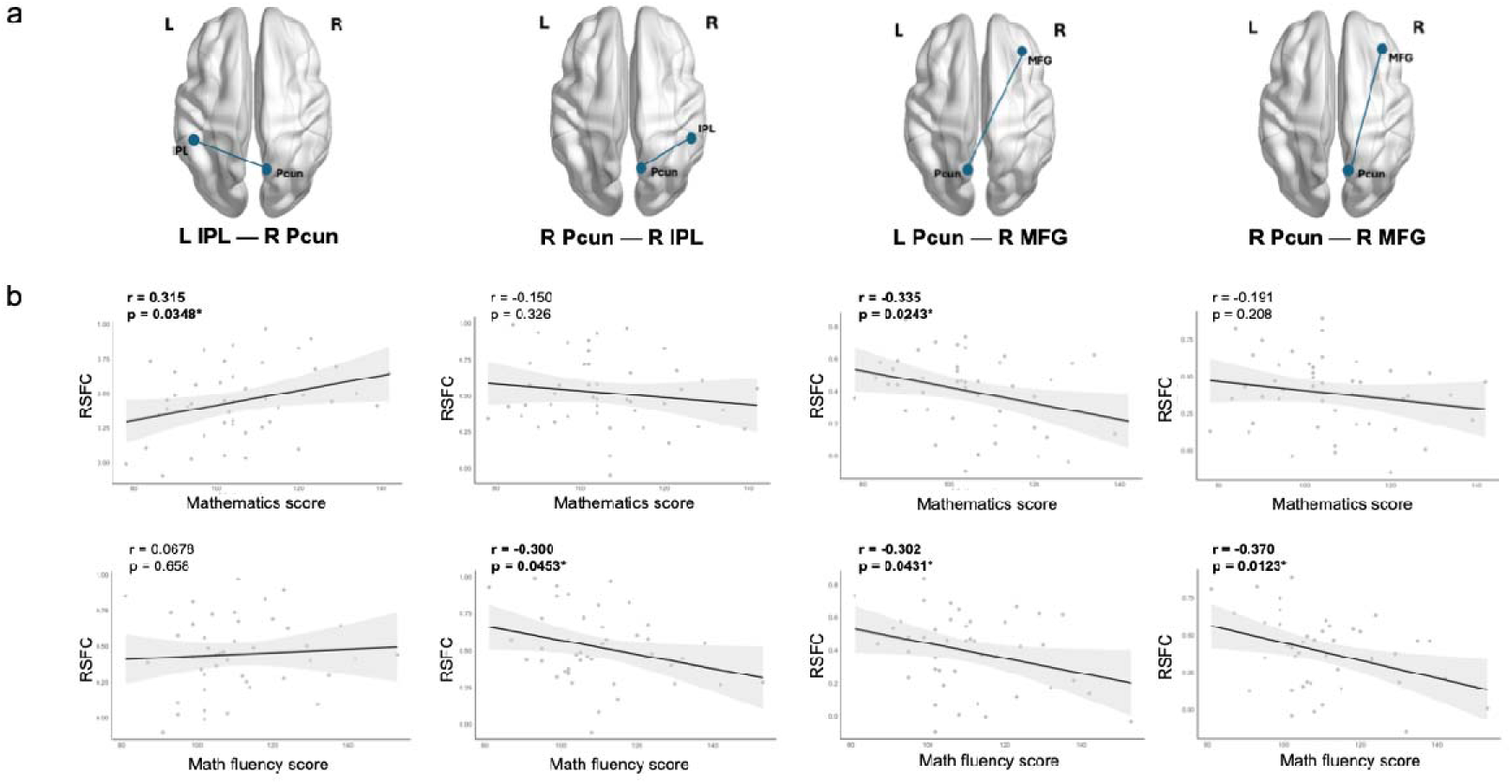
RSFC and significant correlations with Mathematics or Math Fluency. **a,** Smoothed brains (axial) visualised by BrainNet Viewer v1.7 [88], showing functional connection between significant ROIs. **b,** Scatter plots of the correlations with best-fit lines ± 95% confidence intervals (shaded regions). Top: RSFC of the respective ROI pair (in **a**) and mathematics scores. Bottom: RSFC of the respective ROI pair (in **a**) and math fluency scores. The Pearson’s correlation values and respective *p* values are indicated, where *\* p* < .05.

The left inferior parietal lobule—right precuneus RSFC had a positive correlation (r = 0.315, *p* < .05) with mathematics scores, while the right precuneus—right inferior parietal lobule RSFC was negatively correlated (r = −0.300, *p* < .05) with math fluency scores. Similarly, the right precuneus—right middle frontal gyrus had a negative correlation (r = −0.370, *p* < .05) with math fluency scores. Finally, the left precuneus—right middle frontal gyrus RSFC had a negative correlation (r = −0.302, *p* < .05) with both mathematics and math fluency scores. (Fig. 3)

#### Validation

Shapiro-Wilk’s test on the four RSFC variables and both mathematics and math fluency, indicated no violation of the normality assumption. (Appendix C.1) Bootstrapping with replacement was used to check for sensitivity of the significant correlation results. Given that none of the confidence intervals of the correlation coefficients included 0, this suggests that the respective variables were most likely truly correlated. (Appendix C.2) The confidence intervals also provide further validation of the correlation directions in Fig. 3.

### 3.3 Comparing RSFC between mathematical competencies

Children were categorised as lower performers (LP) or higher performers (HP), based on the 10^th^ percentile scores (Table 3).

**Table 3.** 10^th^ percentile scores of Mathematics and Math Fluency used to classify LP in each domain.

| Domain | N | Score |  |  | N (LP) |
| --- | --- | --- | --- | --- | --- |
|  |  | Maximum | Minimum | 10<sup>th</sup> Percentile |  |
| Mathematics | 45 | 142 | 78 | 87.4 | 5 |
| Math Fluency | 45 | 153 | 81 | 95.0 | 7 |

The right precuneus—left inferior parietal lobule RSFC was significantly different between HP and LP in mathematics (Table 4). This aligns with the correlation results in Fig. 3, suggesting that arithmetic problem-solving may be supported by the RSFC between these regions. Considering the potential lack of power due to the small sample sizes used, Cohen’s d was computed for each test to assess effect sizes. Although the RSFC difference between HP and LP in the left precuneus—right middle frontal gyrus was non-significant, the modest effect size (Table 4) nonetheless hints at likely differences, in line with the correlation results (Fig. 3). Importantly, these findings show that the right precuneus—left inferior parietal lobule RSFC may be associated with children’s mathematical competency.

**Table 4.** Independent samples t-test of differences between RSFC of HP vs LP for Mathematics.

| RSFC | HP (N = 40) |  | LP (N = 5) |  | Independent samples t-test |  |  |  |
| --- | --- | --- | --- | --- | --- | --- | --- | --- |
|  | M | SD | M | SD | t | Sig. (2-tailed) | Cohen's d | 95% CI |
| L IPL – R |  |  |  |  |  |  | 1.04 |  |
| Pcun | .47 | .24 | .21 | .33 | 2.20 | .033* | (strong) | [ 0.02, 0.50 ] |
| R Pcun – |  |  |  |  |  |  | 0.04 |  |
| R IPL | .52 | .24 | .51 | .27 | .09 | .929 | (weak) | [ -0.22, 0.24 ] |
| L Pcun – |  |  |  |  |  |  | -0.40 |  |
| R MFG | .39 | .24 | .48 | .09 | -.85 | .400 | (modest) | [ -0.31, 0.13 ] |
| R Pcun – |  |  |  |  |  |  | 0.07 |  |
| R MFG | .39 | .24 | .37 | .29 | .16 | .874 | (weak) | [ -0.21, 0.25 ] |

Note. \* $p < .05$ .

No RSFC pairs with significant correlations to math fluency (Fig. 3) showed a significant difference between HP and LP in math fluency (Table 5). However, effect sizes reveal that RSFC between the right precuneus and both the ipsilateral inferior parietal lobule and middle frontal gyrus, showed relatively stronger differences between HP and LP. These support the correlation results (Fig. 3), highlighting the involvement of these regions in cognitive processes relevant to math fluency tasks. Interestingly, similar correlation values were computed for math fluency and the left precuneus—right middle frontal gyrus RSFC (r = −0.302, p = 0.0431 < .5), and the right precuneus—right inferior parietal lobule RSFC (r = −0.300, p = 0.0453 < .5) (Fig. 3). However, the former had a moderate effect size (d = −0.68), while the latter had a modest effect size (d = −0.46). This suggests that the right precuneus—right inferior parietal lobule RSFC may not be a robust biomarker of math fluency competence, even with larger sample sizes.

**Table 5.** Independent samples t-test of differences between RSFC of HP vs LP for Math Fluency.

| RSFC | HP (N = 38) |  | LP (N = 7) |  | Independent samples t-test |  |  |  |
| --- | --- | --- | --- | --- | --- | --- | --- | --- |
|  | M | SD | M | SD | t | Sig. (2-tailed) | Cohen's d | 95% CI |
| L IPL – R |  |  |  |  |  |  | 0.35 | [ -0.12, |
| Pcun | .46 | .24 | .37 | .37 | .85 | .401 | (modest) | 0.31 ] |
| R Pcun – |  |  |  |  |  |  | -0.68 | [ -0.35, |
| R IPL | .50 | .24 | .66 | .23 | -1.65 | .107 | (moderate) | 0.04 ] |
| L Pcun – |  |  |  |  |  |  | -0.46 | [ -0.29, |
| R MFG | .38 | .24 | .48 | .16 | -1.12 | .267 | (modest) | 0.08 ] |
| R Pcun – |  |  |  |  |  |  | -0.69 | [ -0.36, |
| R MFG | .36 | .23 | .53 | .26 | -1.69 | .099 | (moderate) | 0.03 ] |

#### Validation

The independent samples t-test assumes homogeneity of variance. Levene’s test for both mathematics and math fluency show that there was homogeneity of variances between HP and LP for the ROI pairs (p >0.05, signifying homoscedasticity) (Appendix D.1). Another crucial assumption of the independent samples t-test is the sample means are normally distributed. The Shapiro-Wilk test of normality showed that with the exception of the right precuneus—right inferior parietal lobule RSFC of LP, all other variables were normally distributed (Appendix D.2).

## 4. Discussion

The study explored neural mechanisms underlying mathematics and math fluency in three regions: middle frontal gyrus, inferior parietal lobule and precuneus, with a sample of school-entry children.

Firstly, we predicted stronger connectivity among all three regions with increasing mathematics scores. The results corroborate as there was a positive correlation between the left inferior parietal lobule—right precuneus RSFC and mathematics scores (Fig. 3). However, there was a negative correlation between the left precuneus—right middle frontal gyrus RSFC and mathematics (Fig. 3). This shows that children who performed better in mathematics not only had higher connectivity between the parietal regions, but also a weaker frontoparietal connectivity. This could indicate that the better performing children had begun the neural shift from frontal to parietal regions. As previous studies have also shown, children rely more on posterior parietal systems for problem-solving and mathematical fact retrieval with development [69]. As mathematics involved several cognitive functions such as logical reasoning and visuospatial processing, the positive connectivity may reflect a better integrated network to support the diverse cognitive functions.

Secondly, math fluency did not show stronger connectivity within posterior parietal regions to aid arithmetic fact retrieval. Instead, we observed a negative correlation between math fluency scores and RSFCs in the: (1) right precuneus—right inferior parietal lobule, (2) left precuneus—right middle frontal gyrus, and (3) right precuneus—right middle frontal gyrus. Math fluency is posited to recruit domain-general functions for quick manipulation of numbers in the working memory. Therefore, the negative correlation could stem from excessive RSFC among both frontoparietal and posterior parietal systems, suggesting inefficient brain integration. Furthermore, as math fluency is reliant on basic fact retrieval while the precuneus supports higher-order mathematical processing, perhaps math fluency might have stronger associations with connectivity between the angular gyrus [[62],[63]] and other memory regions like the hippocampus. Future studies may consider the connectivity between these regions.

Thirdly, we also anticipated distinct neural mechanisms for mathematics and math fluency. The hypothesis partially corroborated with the results. With the exception of the left precuneus—right middle frontal gyrus RSFC which showed a negative correlation with both scores, all other correlations were significant between the RSFC pairs and mathematics *or* math fluency. This shows that the neural mechanisms supporting mathematics and math fluency are likely different, despite having inherent similarities.

It is of note that presently, there is no standard for expounding on the practical significance of RSFC between brain regions. The interpretation that higher connectivity is better or worse for mathematical achievement is typically a post-hoc explanation of the correlation being positively or negatively related to mathematics scores [89]. More importantly, there is greater potential in using the results to understand differing competencies in mathematics.

The left inferior parietal lobule—right precuneus RSFC was significantly different in LP children in WIAT-III’s Mathematics subtest, compared to the HP children (Table 4). However, all other RSFC pairs (presenting significant correlations (Fig. 3) with mathematics or math fluency scores) were not significantly different in LP children. While we found differences in the left inferior parietal lobule—right precuneus RSFC in LP children, these regions were not frontoparietal as predicted. The results suggest further integration among posterior parietal brain regions may have occurred after the frontal to parietal shift, consistent with findings from other studies [[57],[90]]. Thus, poorer mathematics performers exhibited different left inferior parietal lobule— right precuneus RSFC compared to better performers, who may have undergone additional strengthening of connectivity in posterior parietal regions. While the frontoparietal shift and development of mathematical skills continue into adulthood [91], crucially, our results imply that connectivity among posterior parietal regions may be more relevant in explaining competency levels than frontoparietal connectivity. Future studies should investigate if RSFC among the posterior parietal regions can better predict mathematical competency compared to frontoparietal regions. Furthermore, as the average age of children in our sample is 6.96 years, this shows that the frontal to parietal shift in activity is likely initiated before age seven [92]. Given the lack of studies confirming the ages during which such connectivity development occurs post-shift, further research should replicate these findings in other samples for better generalisability and reliability.

### 4.1 Implications for education

Firstly, understanding the neural differences between mathematics and math fluency holds significance for education strategies. As math fluency demands the retrieval of basic arithmetic facts, students who struggle with this are naturally hindered in their ability to engage in higher-order mathematical learning. In our study, not all students who performed poorly in mathematics also performed poorly in math fluency. However, studies have demonstrated that brain measures are more predictive of growth in mathematical abilities from childhood than behavioural measures such as assessment scores [53]. Therefore, it is plausible that proficiency in math fluency contributes to proficiency in mathematics (affirming the pyramid model that fluency is the foundation of mathematical reasoning and problem-solving), and mathematics may additionally contribute as an indicator of relative competency. Perhaps fluency provides the basis for reasoning through the understanding of quantities. Since the brain mechanisms supporting mathematics and math fluency are different, it is important for educators to intentionally improve both aspects of mathematics. Secondly, these results demonstrate that children from the same grade and ages may exhibit differing speeds of development neurobiologically, resulting in different mathematical competencies. This provides further evidence of the need for subject banding systems (where children who perform at similar levels are taught together), such that targeted strategies can support children developing at different speeds. This way, children who perform poorer than their peers are not demoralised early into their education, resulting in a vicious cycle of self-doubt and an inability to learn new mathematical skills [93]. This is particularly important in education systems which mark on a bell-curve in examinations and inherently spur competition among peers [94].

A study by Delazer and colleagues [57] showed that in adults, after extensive practice, two training methods, drill (i.e., rote learning) and strategy (i.e., understanding the underlying problem), led to significant improvements in speed and accuracy, although training by strategy led to higher accuracy and better transfer to new problems. Contrasting brain activation during both training methods, differential activation of the precuneus region emerged. As activation in the precuneus is often found in episodic memory retrieval [95], this implies that subjects trained by strategy developed visual imagery strategies to solve new problems, as opposed to subjects in drill learning where problems were solved through memory.

### 4.2 Limitations and future research

An important but inherent limitation of using fNIRS is its spatial resolution may limit the accuracy of the results. For instance, within the inferior parietal lobule, fMRI studies have found dissociable patterns of relations between arithmetic competence in children among subdivisions of the intraparietal sulcus and angular gyrus [96]. Other MRI studies have also found that RSFC patterns in subareas of the precuneus are distinct, which results in different interactions between subregions of the precuneus and other regions (e.g., intraparietal sulcus) [97]. Future studies may examine the feasibility of narrowing down these regions using fNIRS and with children, and investigating if doing this would dispute the findings of this study.

Next, we used a seed-based analysis approach for RSFC. This relies on an initial hypothesis of the regions and does not fairly investigate all brain regions and their potential roles in mathematical proficiency. Therefore, future research could consider mathematical proficiency in children beyond these regions, such as the hippocampus involved in memory. Additionally, factors such as the methods (e.g., parametric or non-parametric), types of variables used, and sample size produce varying statistical results even with identical data[[98],[99],[100]]. Future studies could use the machine learning approach which allows for validation techniques to interpret the findings [[101],[102]]. One example is to classify and predict neuroimaging features of the different neural mechanisms supporting mathematics versus math fluency.

