## Appendix A for "Comparing neurocognitive mechanisms of mathematical ability and fluency in children: insights from an fNIRS study"

Sources, detectors and channels based on the international 10-20 standard system.

| Source(s) | Detector(s) | Channel(s) | Region | Brain Atlas |
|---|---|---|---|---|
| AF7, AF3, F3 | F5, F1 | 5, 8, 14, 15 | Dorsolateral Prefrontal Cortex (BA 46) | Brodmann |
| CP5, P3 | CP3, P5 | 21, 22, 31, 32 | Inferior Parietal Lobule:<br>• Intraparietal Sulcus (BA 39 & 40),<br>• Angular Gyrus (BA 39),<br>• Supramarginal Gyrus (BA 40)*<br>**slightly overlapping region* | Brodmann |
| CP1, Pz | P1 | 25, 35 | Precuneus | AAL2 |
