## Appendix B for "Comparing neurocognitive mechanisms of mathematical ability and fluency in children: insights from an fNIRS study"

Pearson's correlation coefficients for all ROIs and Mathematics or Math Fluency scores.

| RSFC | Mathematics | Math Fluency |
|---|---|---|
| L IPL & L MFG | -.121 | -.0957 |
| L IPL & R MFG | -.100 | -.0676 |
| R IPL & L MFG | -.0573 | -.0852 |
| R IPL & R MFG | -.0678 | -.126 |
| L IPL & L Pcun | -.0620 | -.0922 |
| L IPL & R Pcun | .315 * | .0678 |
| R IPL & L Pcun | -.161 | -.221 |
| R IPL & R Pcun | -.150 | -.300 * |
| L MFG & L Pcun | -.263 | -.189 |
| L MFG & R Pcun | -.147 | -.212 |
| R MFG & L Pcun | -.335 * | -.303 * |
| R MFG & R Pcun | -.191 | -.370 * |
| L IPL & R IPL | .0104 | -.110 |
| L MFG & R MFG | -.0315 | -.0329 |
| L Pcun & R Pcun | -.110 | -.142 |

Note. * $p < .05$.
