## Appendix C for "Comparing neurocognitive mechanisms of mathematical ability and fluency in children: insights from an fNIRS study"

Appendix C.1. Shapiro-Wilk Test of Normality.

| | Shapiro-Wilk Test of Normality | |
|---|---|---|
| Variable | Test statistic | Sig. (2-tailed). |
| Mathematics | .97 | .299 |
| Math Fluency | .95 | .076 |
| L IPL – R Pcun | .98 | .794 |
| R Pcun – R IPL | .97 | .415 |
| L Pcun – R MFG | .97 | .279 |
| R Pcun – R MFG | .99 | .826 |

Appendix C.2. Confidence intervals of significant correlations after bootstrapping.

| | **Bootstrapping *(n = 1000)*** |
|---|---|
| Significant correlations | 95 % CI *(r)* |
| L IPL – R Pcun & Mathematics | [ 0.06, 0.52 ] |
| R Pcun – R IPL & Math Fluency | [ -0.51, -0.07 ] |
| L Pcun – R MFG & Mathematics | [ -0.55, -0.11 ] |
| L Pcun – R MFG & Math Fluency | [ -0.55, -0.00 ] |
| R Pcun – R MFG & Math Fluency | [ -0.59, -0.10 ] |
