## Appendix D for "Comparing neurocognitive mechanisms of mathematical ability and fluency in children: insights from an fNIRS study"

Appendix D.1. Levene's Test between the RSFC of HP and LP for both Mathematics and Math Fluency.

| | | Levene's Test of Homogeneity of Variances | | | |
|---|---|---|---|---|---|
| Domain | RSFC | F-value | df 1 | df 2 | Sig. (2-tailed) |
| Mathematics | L IPL – R Pcun | .48 | 1 | 43 | .492 |
| | R Pcun – R IPL | .36 | 1 | 43 | .552 |
| | L Pcun – R MFG | 3.78 | 1 | 43 | .058 |
| | R Pcun – R MFG | .04 | 1 | 43 | .843 |
| Math Fluency | L IPL – R Pcun | 3.79 | 1 | 43 | .058 |
| | R Pcun – R IPL | .01 | 1 | 43 | .943 |
| | L Pcun – R MFG | 1.77 | 1 | 43 | .190 |
| | R Pcun – R MFG | .21 | 1 | 43 | .652 |

Appendix D.2. Shapiro-Wilk Test of Normality.

| | | | Shapiro-Wilk Test of Normality | |
|---|---|---|---|---|
| Domain | Variable | | Test statistic | Sig. (2-tailed). |
| Mathematics | L IPL – R Pcun | HP | .98 | .737 |
| | | LP | .91 | .483 |
| | R Pcun – R IPL | HP | .98 | .737 |
| | | LP | .69 | .007** |
| | L Pcun – R MFG | HP | .97 | .415 |
| | | LP | .98 | .939 |
| | R Pcun – R MFG | HP | .98 | .748 |
| | | LP | .88 | .315 |
| Math Fluency | L IPL – R Pcun | HP | .98 | .876 |
| | | LP | .93 | .588 |
| | R Pcun – R IPL | HP | .98 | .679 |
| | | LP | .88 | .225 |
| | L Pcun – R MFG | HP | .97 | .357 |
| | | LP | .99 | .993 |
| | R Pcun – R MFG | HP | .98 | .720 |
| | | LP | .93 | .535 |

Note. ** $p < .01$.
